# T cell costimulatory and inhibitory signals differentially modulate LAT condensate nucleation propensity after TCR ligation

**DOI:** 10.1101/2025.11.01.685673

**Authors:** Saehyun Choi, Sungi Kim, Jessica Tijham, Timothy Eisen, Shumpei Morita, Mark K. O’Dair, Jay T. Groves

**Author notes:** These authors are equally contributed (Saehyun Choi, Sungi Kim).

## Abstract

Linker for activation of T cells (LAT) is a membrane-surface scaffold protein that assembles key signaling molecules upon T cell receptor (TCR) activation. Recent observations have revealed LAT assembly occurs in discretized condensates traceable to single molecular TCR activation events. Moreover, the condensation process occurs abruptly, but after a distinct delay, indicative of a type of phase transition. Here we examine the effects of costimulatory (CD80, CD86) and inhibitory (PD-L1) ligands on the nucleation propensity of LAT condensation. We utilize single molecule mobility tracking to resolve the moment a pMHC molecule binds to TCR combined with intracellular imaging to measure the time delay between pMHC:TCR binding and nucleation of the corresponding LAT condensate. These delay time distributions reflect the propensity function for stochastic nucleation of the LAT condensate and provide a measure of the effective signal strength from a single, activated TCR. The results reveal that CD28 stimulation by CD80 or CD86 causes a distinct reduction in the mean delay time to LAT condensation while PD-L1 is mildly antagonistic to the process. Collectively, these observations strengthen evidence that probability and timing of the LAT nucleation process itself is tightly associated with signal propagation downstream from individual TCR activation events.

## Introduction

Tells cells exhibit exquisite sensitivity and selectivity during antigen discrimination, with the ability to respond to even one molecule in a complex cell-cell interface^1–5^. Whereas the basic molecular signaling pathway downstream from TCR activation has been known for many years,^6^ specific molecular mechanisms that enable single molecule sensitivity are only beginning to emerge. A key insight is provided by observations that discrete LAT condensates form in response to individual TCR activation events. These experiments revealed that a single pMHC:TCR complex, even under very dilute ligand densities such that there are no other complexes within microns, is capable of inducing nucleation of a LAT condensate containing hundreds of LAT molecules, along with other scaffold and signaling molecules.

The basic idea of a LAT cluster or signalosome, as it has been called in earlier work, has long been known to be a central feature of TCR signaling^7–10^. Upon pMHC binding to TCR, the Src family kinase Lck phosphorylates tyrosine sites on the ITAM domains, to which the Syk family kinase Zap70 is recruited and subsequently activated.^6^ Zap70 has rather distinct substrate specificity and uniquely phosphorylates LAT. As LAT becomes phosphorylated on multiple tyrosine sites, other molecules including Grb2, SOS1, GADS, SLP76, ITK and PLCg1 are also recruited, forming a signaling complex that mediates outgoing signals to both MAPK and calcium signaling pathways. Classically, once Zap70 is activated at the TCR, that TCR was considered to be signaling active and it was generally imagined that multiple active TCR cooperated to drive further downstream signaling through LAT. Early experiments with very high pMHC ligand densities (~ 100 molecules/µm^2^) readily revealed formation of pMHC:TCR microclusters, which are ultimately driven towards the cell center to form the so-named central supra molecular activating cluster (cSMAC).^11–13^ While it quickly became evident that the cSMAC itself was not required for T cell activation^11–13^, the TCR microscluster as a signaling unit remained a compelling concept for many years^17–20^.

The TCR microcluster concept as an essential signaling unit, however, was always difficult to rectify with the well-established fact that T cells respond to as little as one molecule of agonist pMHC^1–5^. More recent experiments focusing on very dilute pMHC densities (e.g. < 1 molecule/µm^2^) provided clear evidence that a lone pMHC:TCR complex is sufficient to drive formation of a LAT condensates^10,21^ and that each LAT condensate is individually competent to induce a cell-wide calcium spike^22^. These experiments demonstrate that cooperativity between multiple TCR is not necessary, but that a high level of cooperativity within the LAT condensate itself gates signal propagation from a single activated pMHC:TCR complex into discrete quanta of output. Another informative observation in these single molecule studies is that the LAT condensation process follows a distinctive trajectory. After the moment of pMHC:TCR binding, there is a delay of tens of seconds (on average) before an abrupt condensation process during which the LAT condensate rapidly grows to hundreds of LAT molecules. This provided the first direct measurement of signal amplification strength between TCR and LAT. A mutation in LAT that increases the rate of phosphorylation at the essential PLCg1 binding site and alters T cell ligand discrimination properties was also found to modulate the delay time to LAT condensation.^10,23^ This result suggests that kinetic features of condensation phase transition itself may play a role shaping signal propagation downstream from TCR. Here, we further examine this possibility by measuring the effect of costimulatory and inhibitory effects on the nucleation propensity of the LAT condensate.

For the studies described here, we utilize single molecule tracking of pMHC ligands, which reveal pMHC:TCR binding events as distinct changes in pMHC mobility. This experimental strategy robustly resolves the position and timing of each pMHC:TCR complex formation event in live, primary T cells. Delay times between initial pMHC:TCR binding and proximal LAT condensation are readily measured. We characterize the effects on LAT condensation delay time from inclusion of costimulatory (CD80 or CD86) and inhibitory PD-L1 ligands in the phantom antigen presenting cell (APC) surface. CD28 stimulation by CD80 or CD86 causes a distinct reduction in the mean delay time to LAT condensation, while preserving the broad shape of the distribution. This is consistent with the general idea that CD28 costimulation enhances upstream kinase activity^24^, but reveals that the mechanism by which this enhancement impacts signaling is traceable to changes in the nucleation propensity of LAT condensates. We also observe a mild elongation of the delay time by inclusion of PD-L1, which could be the result of increased SHP-2 phosphatase activity at the TCR^25,26^. Additionally. these modulations to LAT condensate nucleation propensity lead to corresponding changes in antigen sensitivity as determined by cell activation measurements tracking translocation of Nuclear Factor of Activated T cells (NFAT). Together, these observations illuminate how upstream modulations of kinase and phosphatase activity by costimulation or inhibitory signals can be translated to downstream signaling through their effects on the nucleation propensity of LAT discrete condensates. Notably, the effects are probabilistic and alter only the propensity or likelihood of condensation. The condensates themselves, corresponding to quanta of signaling activity, are not altered.

## Results

### Measuring LAT condensate nucleation delay times

We investigate LAT condensation in response to individual molecular pMHC:TCR binding events using a hybrid live cell-supported bilayer experimental platform. Experiments are performed using a supported membrane functionalized with the adhesion molecule ICAM-1 (~100 molecules/µm^2^) and low densities of the strong agonist MCC pMHC (0.2~0.5 molecules/µm^2^). Primary CD4+ T cells with the AND TCR form immunological synapses with this phantom APC surface in a hybrid live cell-supported membrane configuration (**Figure 1a**). We additionally functionalize bilayers with inhibitory PD-L1 ligand at ~ 1000 molecules/µm^2^, co-stimulatory CD80 ligand at ~ 1700 molecules/µm^2^ and co-stimulatory CD86 ligand ~2500 molecules/µm^2^ in specific experiments, which provide very high costimulatory or inhibitory signals.^4,27,28^ Fluorescent fusion proteins, including LAT-eGFP and NFAT-mCherry were expressed using the PLAT-E retroviral platform. The NFAT construct used here is a truncated form of NFAT1 containing the regulatory domain involved in the nucleocytoplasmic shuttling of NFAT but lacking the DNA binding domain, which has been validated as a sensor for signaling downstream from calcium in T cells^4,5,10^. For NFAT titration experiments, LAT-eGFP and NFAT-mCherry were co-expressed by using self-cleaving P2A peptide (see SI for experimental details).

**Figure 1.**
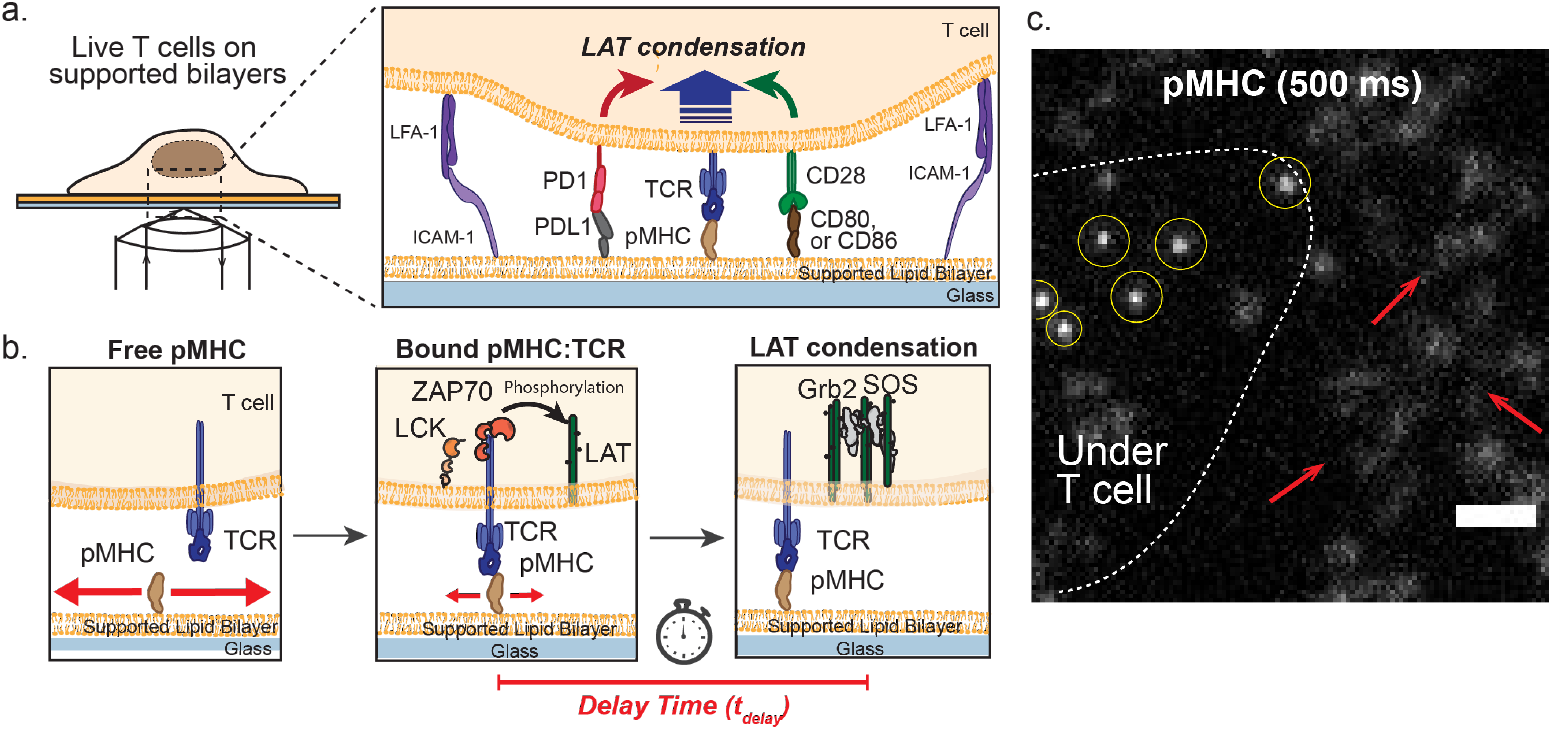
Measurement of single pMHC:TCR binding events and LAT condensation in presence of inhibitory and costimulatory ligands. **(a)** Experiment setup of live-cell T cells imaging on the supported lipid bilayers by TIRF microscopy. Supported lipid bilayers are functionalized with pMHC (0.1-0.3 µm^-2^) in presence or absence of inhibitory (PDL1) or costimulatory (CD80, CD86) ligands. All bilayers contain ICAM-1 ligand (~100 µm^-2^) for T cell adhesion. **(b)** Schematic of T cell signaling that occurs downstream of pMHC:TCR binding. With our single-molecule imaging method, we identify bound pMHC to TCR and observe the following LAT condensation in live T cells. **(c)** pMHC-atto647 image with long exposure (500 ms) showing bound pMHC (yellow circles) and unbound pMHC (red arrows). The white dotted line represents the boundary of cell membrane. Scale bar, 2 µm.

Total Internal Refelection Fluorescence (TIRF) microscopy of the live T cells on supported bilayers enables single-molecule imaging of pMHC:TCR binding events while simultaneously tracking LAT condensate formation^3–5,10^ (**Figure 1b, c**). At low pMHC densities (< ~0.5 molecules/µm^2^), individual pMHC molecules undergoing lateral Brownian motion on the bilayer are individually traceable. Upon binding to a TCR, the pMHC molecule’s mobility is significantly reduced by the ligation event. When pMHC is imaged with a long-exposure time (500 ms), the rapidly moving unbound pMHC are blurred out. In contrast, the slow-moving, bound pMHC:TCR complexes appear as well resolved, high-intensity spots. This method can robustly resolve bound complexes from free ligands with better than 95% certainty^5^. We characterize LAT condensate nucleation propensity by compiling delay times between initial binding of pMHC to TCR and nucleation of the corresponding LAT condensate^10,21^.

Nucleation, growth, and dissolution of LAT condensates are readily resolved in TIRF imaging of live primaty T cells (**Figure 2**). At the low pMHC densities used in these experiments, pMHC:TCR complexes are spaced microns apart and LAT condensates are similarly well isolated in general. This spatial separation facilitates accurate attribution of individual LAT condensates to the originating pMHC:TCR binding events. As shown in **Figure 2a**, we observe productive pMHC:TCR binding events that initiate LAT condensation in their vicinity, alongside unproductive pMHC:TCR binding events that fail to trigger sustained LAT condensation. We note instances, such as event 5 in **Figure 2a, b**, where transient fluctuations in LAT density occur, but fail to develop into a stable condensate. To measure a delay time between pMHC:TCR binding events and the onset of LAT condensation, we track LAT-eGFP intensity in time-series TIRF microscopy images using ImageJ Trackmate. We identify the onset of LAT condensation as the time when sustained accumulation is visible (**Figure 2c**). Notably, the size and lifetimes of LAT condensates do not correlate with pMHC:TCR binding dwell time^10^. Once nucleated, all LAT condensate have similar properties as well as signaling output^22^.

**Figure 2.**
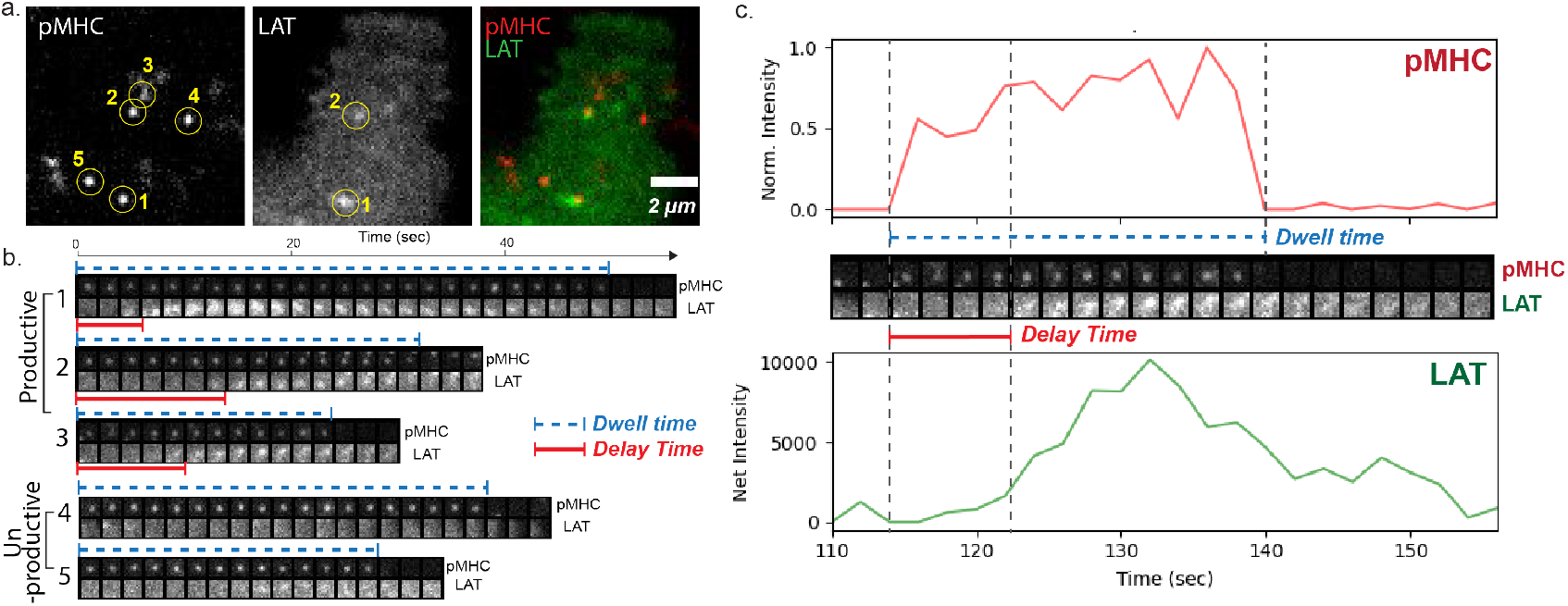
TIRF microscopy images of pMHC on supported bilayers and primary mouse CD^4+^ T cells expressing AND TCR and LAT-eGFP. **(a)** Representative TIRF images of pMHC, LAT-eGFP channels and an overlay of those two channels. pMHC is loaded with fluorescently labeled agonist peptide Atto647-MCC. Five pMHC:TCR binding events are numbered, and only subset of them produces LAT condensates. **(b)** Timelapse images show both productive (event 1, 2, 3) and unproductive (event 4, 5) pMHC:TCR binding events for LAT condensation. Delay time is estimated as the time gap in between the onset of pMHC:TCR binding events and the onset of LAT condensation (noted by the red line). Dwell time of pMHC:TCR binding events are shown in blue dotted line. **(c)** Intensity traces of MCC(Atto647) pMHC and LAT condensates.

### Costimulatory and inhibitory signal effects on LAT condensate nucleation propensity

When looking at LAT condensation as a discrete stochastic process, the propensity function is the momentary kinetic rate (probability per unit time) of a LAT condensation event occurring at a particular delay time from the original pMHC:TCR binding event that triggered it (see McAffee et al^10^ for details). For LAT condensate nucleation in T cells, the propensity function exhibits a complex time-dependence, reflecting kinetic proofreading at the TCR (first few seconds) as well as the stochastic condensate nucleation mechanism itself at longer times^10,21^. Ultimately, the propensity function reflects the momentary state of balance between activated Zap70 kinases on the TCR and locally competing phosphatases. The ensuing LAT phosphorylation-dephosphorylation reactions ultimately set the probability of LAT condensation, which occurs when a localized fluctuation in phosphorylated LAT levels is sufficient to nucleate crosslinking^29,30^. Once nucleated, the LAT condensate grows quickly as positive feedback mechanisms, including phosphatase exclusion^9^ and others, shift the balance in favor of phosphorylation. Perturbations to this balance are reflected an altered propensity function and lead to corresponding changes in antigen sensitivity^10^.

Here we characterize how the LAT condensation process is modulated by CD28 costimulatory or PD-1 inhibitory signals. We first confirm that pMHC:TCR binding kinetics themselves are not influenced by costimulatory or inhibitory signals. Measured distributions of *in situ* pMHC:TCR binding dwell times are shown (grey bars) for each condition studied in **Figure 3**. The dashed line fits correspond to first order dissociation kinetics along with a static parameter (plateau value at long times), which accounts for rare, extremely long-lived features (possibly pMHC aggregates). Measured dissociation rate constants for the first order process were consistent around 0.1 s^-1^, which is in line with typical values for the strong MCC pMHC interaction with AND TCR. Also indicated on the plots are the subset of binding events that triggered LAT condensation overlaid (colored bars). Collectively, these measurements of all pMHC:TCR binding events along with identification of which ones successfully nucleate LAT condensates amount to mappings of the LAT condensate nucleation propensity function. This can be seen in the rising success probability over the first 20 s followed by a stabilization or decay in success rate at later delay times after. All conditions examined follow this pattern, indicating that the basic LAT condensation mechanism is essentially preserved. We also note that both the size of LAT condensates are unaltered by CD28 costimulatory or PD-1 inhibitory signals (**Figure S1**).

**Figure 3.**
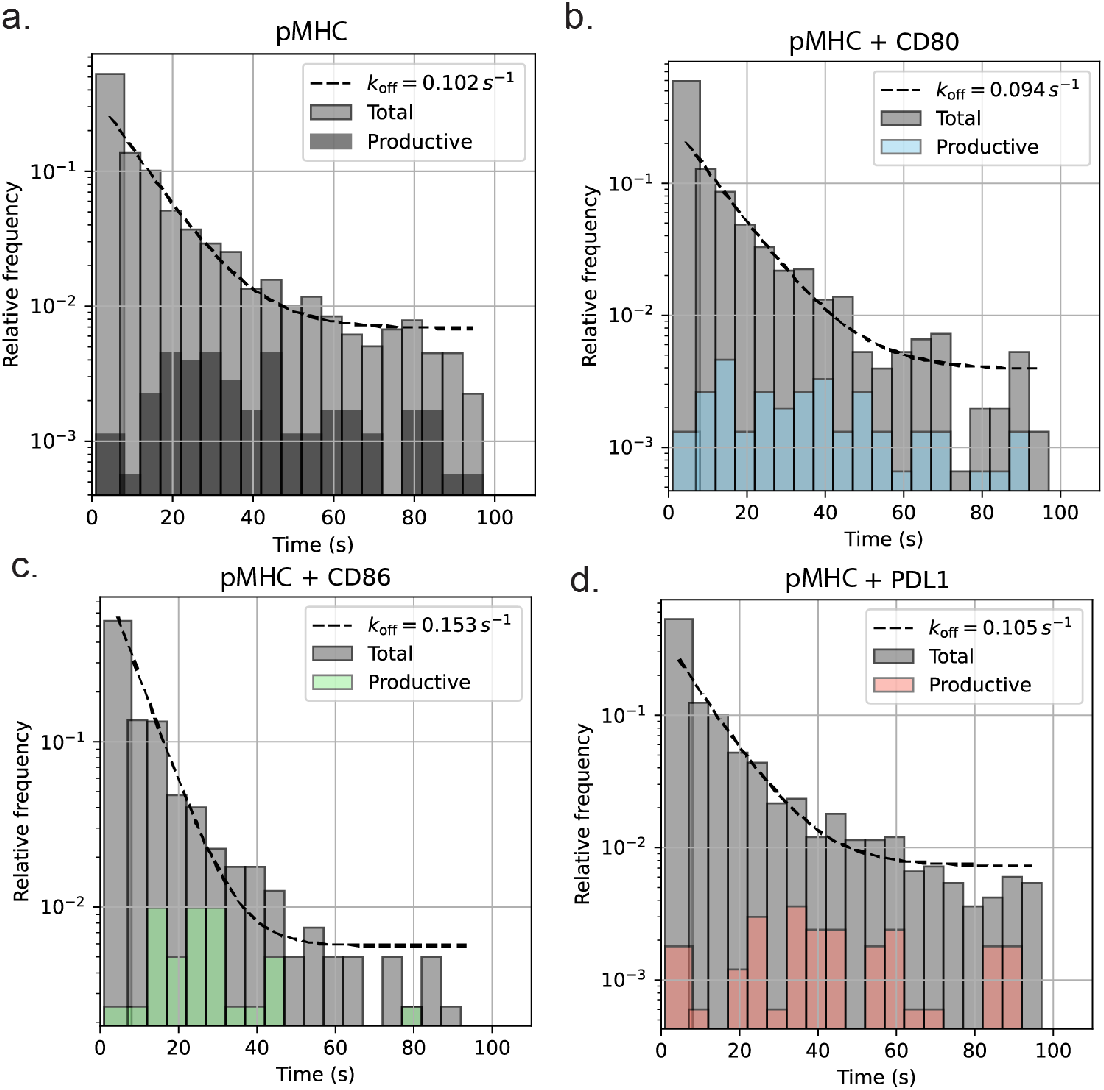
Co-stimulatory or co-inhibitory do not alter pMHC:TCR binding kinetics significantly. T cells are imaged on supported lipid bilayers contain **(a)** pMHC alone, **(b)** pMHC with CD80, **(c)** pMHC with CD86, and **(d)** pMHC with PDL1. pMHC is loaded with fluorescently labeled agonist peptide Atto647-MCC. All bilayers are functionalized with ICAM ligand for cell adhesion. pMHC:TCR dwell time distributions are measured by tracking single-molecule tracks of atto647-MCC that are under T cells. Data points were collected from sampling as following: MCC (n = 28 cells, 1277 events; 67 with LAT condensation), MCC+PDL1 (n = 16 cells, 1135 events; 40 with LAT condensation), MCC+CD80 (n = 18 cells, 1092 events; 47 with LAT condensation), and MCC+CD86(14 cells, 400 events; 21 with LAT condensation). “Productive” distribution indicates a subset of pMHC:TCR binding events that successfully induced LAT condensation.

Next, we examine how the delay time from pMHC:TCR binding events to LAT condensation are influenced by costimulatory and inhibitory signals. Although full pathways of CD28 and PD1 receptor signaling are multifaceted^31–33^, they both have potential to modulate upstream, TCR proximal kinase-phosphatase balance^24,31^. We observe that CD28 stimulation (via CD80 or CD86) caused a distinct reduction in the mean delay time to LAT condensation, relative to agonist pMHC and ICAM-1 alone (**Figure 4 a-c**). In contrast, PD-1 stimulation by PD-L1 produced a modest elongation in the delay time (**Figure 4d)**.

**Figure 4.**
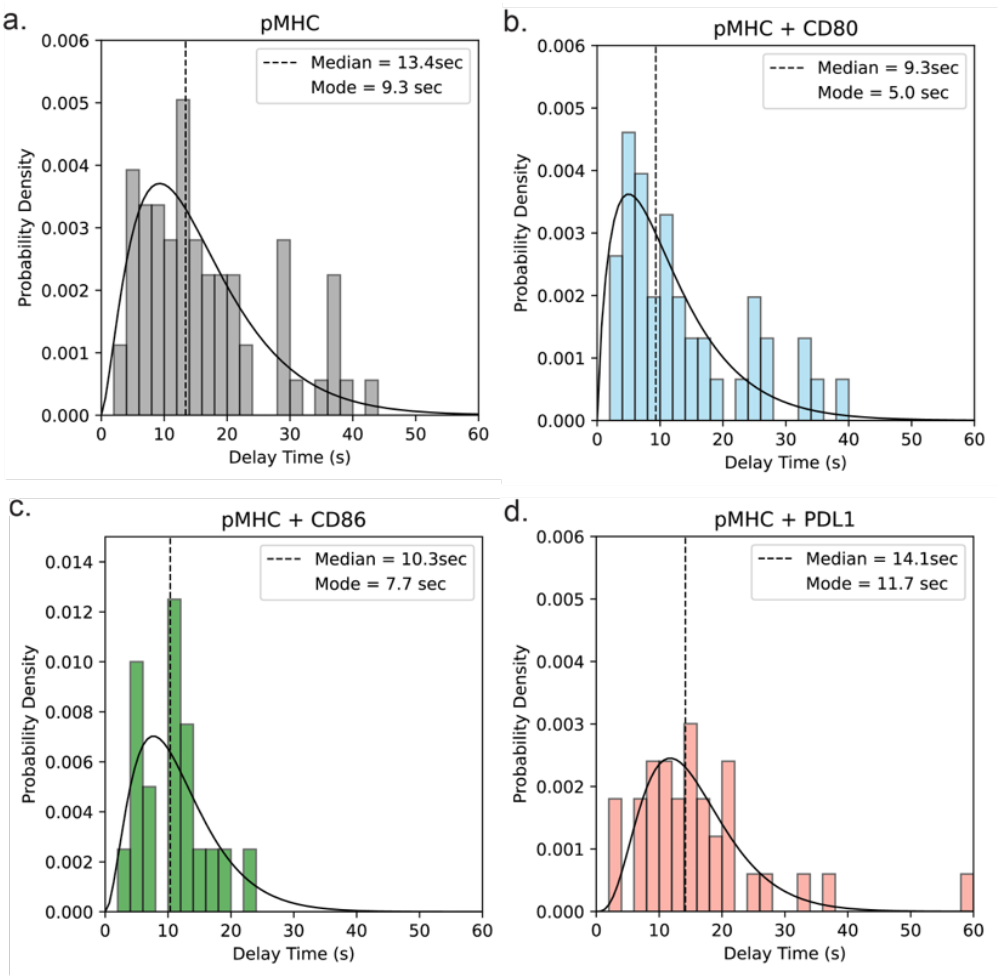
The delay time between pMHC:TCR binding events to LAT condensation in presence of co-inhibitory and co-stimulatory ligands. All supported lipid bilayers functionalized with pMHC loaded with agonist peptide MCC-Atto647 and ICAM-1. **(a)** pMHC alone, **(b)** pMHC with PDL1, **(c)** pMHC with CD80, and **(d)** pMHC with CD86. Both median and mode of distribution shows the consistent trends of altered delay time from pMHC:TCR binding events to LAT condensation in presence of co-stimulatory and co-inhibitory signals.

### CD28 costimulation selectively enhances T cell responsiveness to weak peptide antigens

Lastly, we assess T cell activation responses under costimulatory, inhibitory, and control conditions. Extracellular calcium influx is one of the earliest cell-wide signals in TCR activation. Recent observations have further revealed that each LAT condensate is individually competent to trigger a singular calcium spike, within seconds of nucleation, to the overall cellular calcium level.^22^ Sufficient calcium activity leads to T cell activation, including cytokine secretion, and translocation of NFAT from the cytosol to the nucleus serves as a robust, cell-wide readout of this transition^4,5,34^. NFAT-mCherry in cytosol was imaged using epi fluorescence microscopy at a plane 3 – 5 µm above the TIRF image plane. NFAT translocation was quantified by measuring the cytosol-to-nucleus intensity ratio in epifluorescence microscopy images (**Figure 5a**).

**Figure 5.**
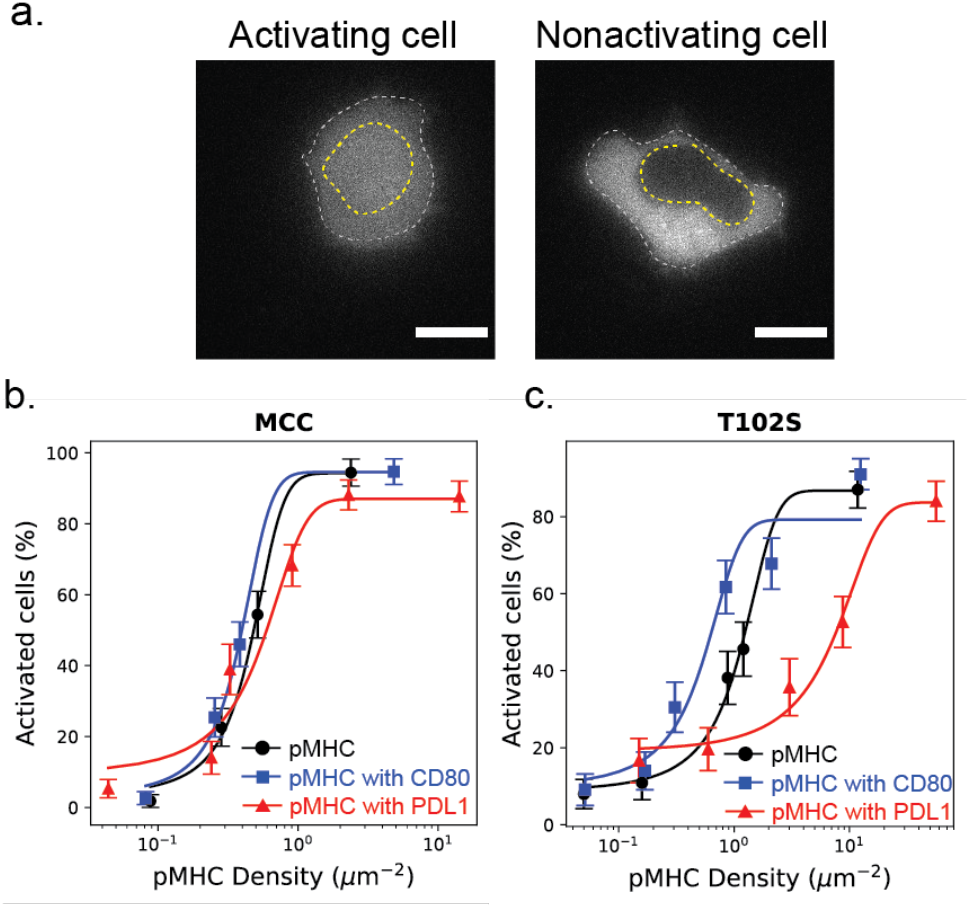
NFAT translocation titration. **(a)** Epi-fluorescent images of NFAT-mCherry transduced T cells interacting with supported lipid bilayers. Analysis of NFAT activation determined by the ratio of mean nuclear intensity (yellow line) to mean cytosolic intensity (white line). Activating cells show brighter nucleus, while nonactivating cells show dark nucleus. Scale bar, 5 μm. (b, c) Density-dependent change in NFAT activation for a population of AND TCR T cells stimulated by pMHC on bilayers that loaded with **(b)** MCC agonist peptide and **(c)** weaker agonist T102S peptide. Each curves represent each condition of pMHC only (blue), pMHC with PDL1 (red), and pMHC with CD80 (blue). Data are from >30 cells per condition.

In the following experiments, we systematically varied the pMHC densities and determined the percentage of activated cells after 10 ~ 30 mins by monitored NFAT translocation. Titration data for stimulation with the strong agonist MCC pMHC, **Figure 5b**) reveal equivalent EC_50_ values of ~0.2 molecules/µm^2^ under all conditions. This result is not surprising given that the overall probability of LAT condensation was similar in all cases as well (e.g. see **Figure 3**). The most notable difference in LAT condensation propensity was measured for CD28 costimulation, and this was an increase in nucleation kinetics rather than a change to overall probability of nucleation with long-binding pMHC. Based on these facts, we hypothesized that costimulation or inhibitory signals would produce larger effects on T cell activation under stimulation by weak agonist pMHC with shorter pMHC:TCR binding lifetimes. To test this, we performed titration experiments with the T102S pMHC, which has a substantially faster kinetic off-rate for unbinding from the AND TCR (binding half-life measurements of MCC and T102S peptides displayed in soluble mouse class II MHC molecule E^k^ binding AND TCR were 11 s and 2s, respectively^35^). The activation response of T cells on T102S pMHC bilayers exhibited significant modulation with costimulatory and inhibitory signals (**Figure 5c**). Relative to the T102S pMHC and ICAM-1 control, we measured a 10-fold increase in EC_50_ for PD-1 stimulation and a 5-fold decrease in the EC_50_ with CD28 costimulation (by CD80).

## Discussion and Conclusion

The data presented here offer a mechanistic framework for understanding how T cells integrate costimulatory (CD28/CD80 or CD28/CD86) and inhibitory (PD1/PDL1) signals. Classical studies have typically focused on signal output, such as calcium, transcription factors, or cytokine release. Our work here reveals that distinct effects of these co-signals are also measurable on the kinetics LAT condensate nucleation—without otherwise changing properties of the condensates. Shifts in the kinase-phosphatase balance proximal to an activated TCR alter the propensity of LAT condensate nucleation, leading to corresponding changes in delay time distribution between pMHC:TCR binding and formation of the corresponding LAT condensate. If pMHC disengages from TCR prior to LAT condensate formation, then no signal is transduced^10,22,29^. In this way, changes to the delay time distribution to LAT condensation are translated into altered sensitivity to pMHC ligands with different TCR binding kinetics. Weaker ligands (faster dissociation kinetics) have a lower probability of triggering a LAT condensate because they have a higher probability of disengaging before condensate nucleation. Weak ligands do not produce weaker LAT condensates. All LAT condensates appear to signal equivalently—defining a preserved quanta of signal information—only their probability of formation is altered. We suggest that the signal digitization achieved through discretized LAT condensation events is key to the T cell’s ability to achieve single molecule sensitivity to antigen pMHC. Weak ligands as well as modulatory signals change only the probability of LAT condensate formation, without changing the signal strength from successful condensation events. This assures that even weak signals maintain a high signal-to-noise ratio above background molecular activity within the cell.

## Acknowledgments

We thank F. Marangoni (Harvard Medical School) for providing the NFAT reporter plasmid; L. Teyton (Scripps Research Institute) and M. Davis (Stanford University) for providing the MHC and ICAM-1 bacmids.

## Author contributions

Conceptualization: J.T.G., S.K. and S.C.; formal analysis: S.C., S.K., and J.T.; investigation: S.C., S.K., J.T., S.M., T.E., and M.K.O.; resources: S.C., S.K., J.T., S.M., T.E., and M.K.O.; data curation: S.C., S.M., and S.K.; writing—original draft: S.C. and J.T.G; writing—review and editing: S.C., S.K., and J.T.G; supervision: J.T.G.; funding acquisition: J.T.G.

## Data and materials availability

The data and materials newly created in this study are available from the corresponding author (JTG) upon reasonable request.

## Supplementary Materials

Materials and Methods Figure SI 1

