## Supplementary Information for "T cell costimulatory and inhibitory signals differentially modulate LAT condensate nucleation propensity after TCR ligation"

***Method***

**T cell culture and transduction**

The mice used in this study were 6- to 20-week-old and the cross of (B10.Cg-Tg(TcrAND)53Hed/J)x(B10.BR-H2k2 H2-T18a/SgSnj) from Jackson Laboratory. All animal work was performed with prior approval by the IACUC committee, Lawrence Berkeley National Laboratory Animal Welfare and Research Committee, under the approved protocol 17703. Murine T cells were harvested from lymph nodes and spleens and stimulated with 1 µM MCC peptide in vitro. Both male and female mice were used. The T cell blasts were treated with IL-2 from second to fifth day after harvest. IL-2 treated T cell blasts were transduced using retroviruses produced in Platinum-Eco cell-derived supernatants. 1

**DNA construct (LAT_eGFP, LAT_eGFP_P2A_NFAT_mCherry)**

The murine stem cell virus (MSCV) backbone was used (similar to AddGene #91975) for retroviral transduction of primary murine T cells. Plasmids were constructed using the Gibson Assembly method and cloned in *E. coli* XL1-BLUE. MSCV-LAT-eGFP was constructed from a plasmid containing cDNA for for *Mus Musculus* Linker for Activation of T cells (NM_010689.3). MSCV-NFAT-mCherry was constructed from pMSCV-NFAT1(1–460)-GFP, a gift from F. Marangoni (Harvard Medical School). We used the truncated form of NFAT1 contains the regulatory domain involved in the nucleocytoplasmic shuttling of NFAT but lacks the DNA binding domain. MSCV-LAT-eGFP_P2A_NFAT_mCherry was subcloned from the constructs above with the addition of the P2A sequence that was ordered as an oligonucleotide from Elim Biopharmaceuticals, Hayward, CA.

**Bilayer preparation**

Supported lipid bilayer membranes were prepared in imaging chambers and then functionalized with mouse ICAM-1 and peptide-loaded MHC (pMHC), as described in our previous works.[1,2] In brief, pMHC was prepared by incubating MCC (ANERADLIAYLKQATK)-GGSC-Atto647N with MHC II I-$E^k$ at 37 °C in peptide loading buffer (PLB-1% (w/v) bovine serum albumin (BSA) in PBS (pH 4.5) with citric acid), 18 to 24 h before imaging. Supported bilayers were prepared on etched cover glasses using small unilamellar vesicles containing the following components: 98 mol% 1,2-dioleoyl-sn-glycero-3-phosphocholine (DOPC, Avanti Polar Lipids) and 2 mol% 1,2-dioleoyl-sn-glycero-3-[(N-(5-amino-1-arboxypentyl)iminodiacetic acid)succinyl] nickel salt (DGS-NTA, Avanti Polar Lipids). ICAM-1 (10 nM, estimated density ~100 $\mu m^{-2}$) and pMHC (concentration adjusted, 1-6 pM for 0.1-0.6 $\mu m^{-2}$) were then added to the chambers and incubated for 30-40 min, then rinsed by imaging buffer (20 mM HBS, 137 mM NaCl, 5 mM KCl, 1 mM CaCl2, 2 mM MgCl2, 1 % BSA, 1 mM glucose at pH 7.5).

**Imaging**

All imaging experiments were performed on a motorized inverted microscope (Nikon Eclipse Ti-E; Technical Instruments, Burlingame, CA) with a motorized Epi/TIRF illuminator, and a motorized stage (MS-2000; Applied Scientific Instrumentation, Eugene, OR). A laser launch with 488-, 560-, and 640-nm diode lasers (Coherent OBIS, Santa Clara, CA) was aligned into a custom-built fiber launch (SolamereTechnology Group Inc., Salt Lake City, UT). For TIRF imaging, laser illumination was reflected through the appropriate dichroic beam splitter (ZT488/647rpc, Z561rdc with ET575LP) to the objective lens [Nikon (1.47, numerical aperture; 100×), TIRF; Technical Instruments, Burlingame, CA]. RICM and epifluorescent excitation were filtered through a 50/50 beam splitter or band-pass filters (D546/10×, ET470/40×, ET545/30×, and ET620/60×). All emissions were collected through the appropriate emission filters (ET525/50M, ET600/50M, and ET700/75M) and captured on an EM-CCD (iXon 897DU; AndorInc., South Windsor, CT). All filters were from Chroma Technology Corp. (Bellows Falls, VT). All microscope hardware was controlled using MicroManager.

**pMHC:TCR binding well time and LAT condensation delay time analysis**

Imaging analysis was conducted using the standard TrackMate plugin[3] for ImageJ[4] in conjunction with a custom python code script to generate image tiles as in **Figure 2**. pMHC:TCR binding events were initially tracked via TrackMate's automated workflows, and then each track was manually refined for accurate linking and detection. The delay time between initial pMHC:TCR binding and nucleation of LAT condensates was determined by identifying the first visibly sustained accumulation of LAT in a growth of condensates.

**NFAT translocation titration**

We fixed the cells 10-30 mins after the cell injection onto pMHC functionalized supported lipid bilayers. NFAT-mCherry in cells were imaged using epi fluorescence microscopy at 3 – 5 µm above supported-lipid bilayers. We then analyze the intensity ratio of NFAT-mCherry between cytosol and nucleus to determine the activated cells, which have the intensity ratio above the threshold of 1.

### Supplementary Figure

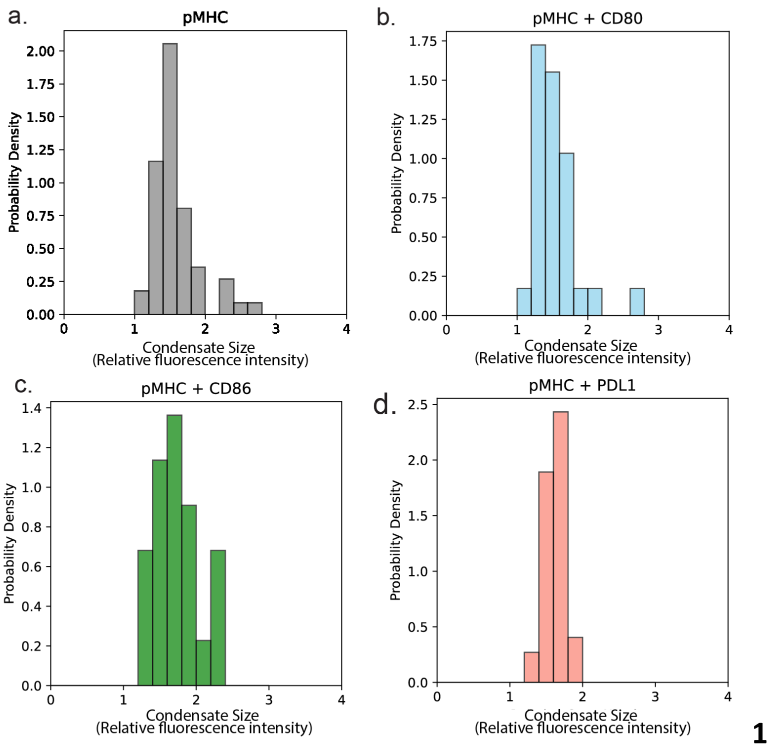


**Figure S1.** LAT condensate size does not change significantly in different conditions. The X-axis indicates the relative protein copy number of LAT, reported as the fold increase compared to the background LAT copy number. The average copy number of LAT in condensates is approximately 250 LAT molecules.[1]

### Reference.


(1) McAffee, D. B.; O'Dair, M. K.; Lin, J. J.; Low-Nam, S. T.; Wilhelm, K. B.; Kim, S.; Morita, S.; Groves, J. T. Discrete LAT Condensates Encode Antigen Information from Single pMHC:TCR Binding Events. *Nat Commun* **2022**, *13* (1), 7446. https://doi.org/10.1038/s41467-022-35093-9.

(2) Lin, J. J. Y.; Low-Nam, S. T.; Alfieri, K. N.; McAffee, D. B.; Fay, N. C.; Groves, J. T. Mapping the Stochastic Sequence of Individual Ligand-Receptor Binding Events to Cellular Activation: T Cells Act on the Rare Events. *Sci. Signal.* **2019**, *12* (564), eaat8715. https://doi.org/10.1126/scisignal.aat8715.

(3) Tinevez, J.-Y.; Perry, N.; Schindelin, J.; Hoopes, G. M.; Reynolds, G. D.; Laplantine, E.; Bednarek, S. Y.; Shorte, S. L.; Eliceiri, K. W. TrackMate: An Open and Extensible Platform for Single-Particle Tracking. *Methods* **2017**, *115*, 80–90. https://doi.org/10.1016/j.ymeth.2016.09.016.

(4) Schindelin, J.; Arganda-Carreras, I.; Frise, E.; Kaynig, V.; Longair, M.; Pietzsch, T.; Preibisch, S.; Rueden, C.; Saalfeld, S.; Schmid, B. Fiji: An Open-Source Platform for Biological-Image Analysis. *Nat. Methods* **2012**, *9* (7), 676–682. https://doi.org/10.1038/nmeth.2019.
